# The Gag Protein PEG10 Binds to RNA and Regulates Trophoblast Stem Cell Lineage Specification

**DOI:** 10.1101/572016

**Authors:** Mona Abed, Erik Verschueren, Hanna Budayeva, Peter Liu, Donald S. Kirkpatrick, Rohit Reja, Sarah K. Kummerfeld, Joshua D. Webster, Sarah Gierke, Mike Reichelt, Keith R. Anderson, Robert J Newman, Merone Roose-Girma, Zora Modrusan, Hazal Pektas, Emin Maltepe, Kim Newton, Vishva M. Dixit

## Abstract

*Peg10* (paternally expressed gene 10) is an imprinted gene that is essential for placental development. It is thought to derive from a Ty3-gyspy LTR (long terminal repeat) retrotransposon and retains Gag and Pol-like domains. Here we show that the Gag domain of PEG10 can promote vesicle budding similar to the HIV p24 Gag protein. Expressed in a subset of mouse endocrine organs in addition to the placenta, PEG10 was identified as a substrate of the deubiquitinating enzyme USP9X. Consistent with PEG10 having a critical role in placental development, PEG10-deficient trophoblast stem cells (TSCs) exhibited impaired differentiation into placental lineages. PEG10 expressed in wild-type, differentiating TSCs was bound to many cellular RNAs including *Hbegf* (Heparin-binding EGF-like growth factor), which is known to play an important role in placentation. Expression of *Hbegf* was reduced in PEG10-deficient TSCs suggesting that PEG10 might bind to and stabilize RNAs that are critical for normal placental development.

## INTRODUCTION

Transposable elements (TEs) are one of the biggest threats to the integrity of prokaryotic and eukaryotic genomes because their insertion into coding or regulatory regions could disrupt essential genes (1-3). Therefore, TEs are often inactivated through mutagenesis (4) or silenced through methylation (5). Some TEs, however, have been repurposed during evolution for the benefit of the host in a process termed domestication (6), and have important roles in development and immunity (7-10).

*Peg10* is a domesticated TE that is expressed in eutherian mammals (8, 11). It has lost the ability to transpose, but retains the retroviral characteristic of frameshifting (FS) that allows the translation of two overlapping reading frames from the same transcript (12). Thus, *Peg10* encodes PEG10-RF1 corresponding to the structural Gag-like protein, as well as PEG10-RF1/2 representing a fusion of the Gag and Pol domains (12). *Peg10*-deficient mouse embryos are not viable because *Peg10* is required for formation of the labyrinth and spongiotrophoblast layers of the placenta (13). Precisely how the PEG10 proteins exert this function is unclear. Whether *PEG10*, which is also expressed in the human testis and ovary (14), has functions outside of the placenta is likewise unclear. Interestingly, *PEG10* is aberrantly expressed in some human tumors including hepatocellular carcinomas (14) and neuroendocrine prostate cancers (15).

Here we study the role of PEG10 in mouse embryonic stem cells (ESCs) and trophoblast stem cells (TSCs) after identifying it as a substrate of the deubiquitinating enzyme USP9X. Expressed in several stem cell populations (16-18) as well as in differentiated cell types, USP9X is essential for mouse development (19). Diverse substrates of USP9X have been reported, including the pro-survival protein MCL-1 (20), the kinase ZAP70 (21), and the E3 ubiquitin ligases SMURF1 (22) and FBW7 (23), but it is unclear if these are critical substrates of USP9X in ESCs and TSCs.

## MATERIALS AND METHODS

### ESCs and TSCs

The Genentech institutional animal care and use committee approved all mouse protocols. *Usp9x*^*fl/y*^ *Rosa26*^*+/CreERT2*^ ESCs were derived from blastocysts after crossing *Rosa26*^*CreERT2*^ (24) and *Usp9x*^*fl*^ (19) mice. Blastocysts were cultured on irradiated mouse embryo fibroblasts (PMEF-N, Millipore Sigma) in ESGRO-2i medium (Millipore Sigma SF016-100) for 13 days to produce ESCs. The cells were subsequently maintained on PMEF-N cells in KNOCK-OUT medium (Invitrogen) supplemented with 1000 U LIF (EMD Millipore), 15% fetal bovine serum (GE Health), 1× non-essential amino acids (Gibco), 2 mM L-glutamine (Gibco), and 50 µM 2-mercaptoethanol (EMD Millipore). A 3xFLAG sequence was inserted after the initiator ATG in *Peg10* by homologous recombination. *Usp9x*^*3xf/y*^ and *Usp9x*^*C1566A/y*^ ESCs were generated by Taconic (Germany) using C57BL/6 NTac ESCs. In brief, a 3xFLAG sequence was inserted after the initiator ATG in *Usp9x* exon 2 (*Usp9x*^*3xf/y*^ ESCs) or the TGT codon encoding Cys1566 in *Usp9x* exon 31 was changed to GCC (*Usp9x*^*C1566A/y*^ ESCs).

TSCs were prepared as described (25) and cultured on plates pre-treated with CellStart substrate (A1014201, Thermo Fisher) for 2 h at 37°C. TSC culture medium was RPMI1670 containing 20% FBS, 1 mM sodium pyruvate, 2 mM L-glutamine, 50 µg/mL penicillin/streptomycin, 100 µM 2-mercaptoethanol, 0.1 mM FGF4 (F2278, Sigma), and 1 µg/mL Heparin (H3149, Sigma). For TSC differentiation, cells were split to a 30% density without CellStart, FGF4 or heparin, and cultured at 20% or 2% oxygen for 5-10 days while changing media every other day. For TSC transfection, cells were plated at 20% density on CellStart with FGF4 and Heparin, and then transfected the day after using Effectene transfection reagent (301425, Qiagen).

PEG10-deficient ESCs and TSCs were generated using a two cut CRISPR strategy. Cells were co-transfected with pRK-Cas9 (38), *Peg10* 5’ sgRNA (CTC TCA CCG CAG CCA TGG C) and 3’ sgRNA (GCA TCA TCC TGC AGT GCT G). sgRNAs were expressed from U6 promoters on individual plasmids.

### Antibodies

Antibodies recognized GAPDH (G9545, Sigma), PEG10 (Genentech monoclonal antibodies were raised against mouse PEG10 residues A2-V377; rat clone 17D4D6C2 was used for western blotting and immunofluorescence, mouse clone 36E2B2B4 was used for immunoprecipitations, and rat clone 5H7B1E3 was used for immuno-gold electron microscopy), USP9X (5679, Genentech), Syntaxin 8 (12206-1-AP, Proteintech), VTI1B (ab184170 Abcam), ABCD3 (ab3421, Abcam), ERK (9102, Cell Signaling), phospho-ERK (9101, Cell Signaling), MEK (4694, Cell Signaling), phospho-MEK (9154, Cell Signaling), p70 S6 Kinase (2708, Cell Signaling), phospho-p70 S6 Kinase (9234, Cell Signaling), Akt (4691, Cell Signaling), phospho-Akt (4060, Cell Signaling), FLAG (A8592, Sigma), HIV1 p24 (ab9071 Abcam), Sall4 (sc-101147, Santa Cruz Biotechnology), and TSG101 (GTX70255, GeneTex).

### Immunoprecipitation and fractionation

Cells were dounced in 10 mM Tris pH 7.4, 7.5 mM KCl, 1.5 mM MgCl_2_, 5 mM 2-mercaptoethanol, plus complete protease inhibitor and PhosSTOP phosphatase inhibitor cocktails (Roche). Insoluble material was removed by centrifugation. The soluble material was adjusted to 20 mM Tris pH 7.4, 135 mM KCl, 1.5 mM MgCl_2_, 2 mM EDTA, 2% Triton, and 20% glycerol before immunoprecipitation of PEG10 or USP9X. Immunoprecipitations with anti-FLAG M2 beads (Sigma) were eluted with FLAG or 3xFLAG peptide as appropriate. Cell lysates were fractionated with NE-PER Nuclear and Cytoplasmic Extraction Reagent (78833, Thermo Scientific).

### Affinity purification mass spectrometry experiments

Protein bands were excised into 10 gel pieces from top to bottom for each pulldown. The pieces were further de-stained in 50 mM NH_4_HCO_3_/50% acetonitrile (ACN), dehydrated in 100% ACN, and re-hydrated in 10 ng/µL trypsin on ice for 20 min. After removing the excess trypsin solution, digestion was performed at 37°C overnight in 25 mM NH_4_HCO_3_. Peptides were extracted in 1% formic acid (FA)/50% ACN, then 100% ACN, and dried in a SpeedVac. After reconstitution in solvent A (2% ACN/0.1% FA), peptides underwent reverse phase chromatography on a NanoAcquity UPLC system (Waters). Peptides were loaded onto a Symmetry® C18 column (1.7 mm BEH-130, 0.1 × 100 mm) with a flow rate of 1 µL/min and a gradient of 2% to 25% Solvent B (0.1% FA/2% water/ACN). Peptides were eluted directly into an Advance CaptiveSpray ionization source (Michrom BioResources/Bruker) with a spray voltage of 1.2 kV, and analyzed using an LTQ Orbitrap Elite mass spectrometer (ThermoFisher). Precursor ions were analyzed in the FTMS at 60,000 resolution; MS/MS data was acquired in the LTQ with the instrument operated in data-dependent mode, whereby the top 15 most abundant ions were subjected to fragmentation.

MS/MS spectra were searched using the Mascot algorithm (Matrix Sciences) against a concatenated target-decoy database comprised of the UniProt Mus musculus protein sequences (downloaded June 2016), known contaminants, and the reversed versions of each sequence. A 50 ppm precursor ion mass tolerance and 0.8 Da fragment ion tolerant were selected with tryptic specificity up to 3 missed cleavages. Fixed modifications were allowed for carbamidomethylated cysteine residues (+57.0215 Da) and variable modifications were permitted for methionine oxidation (+15.9949 Da). Peptide assignments were first filtered to a 1% False Discovery Rate (FDR) at the peptide level and subsequently to a 2% FDR at the protein level. Peptide Spectral Matches (PSMs) per protein were summed across all fractions from the GelC-MS experiment for each IP separately. SAINTExpress-spc v.3.6.1 (26) was run with default settings, comparing the sum of PSMs for all identified proteins enriched with the IP antibody to their negative controls, for each AP-MS experiment separately. Interactions with a SAINT score > 0.9, Bayesian False Discovery Rate (BFDR) < 0.05, and average sum PSM count > 10 were marked.

### Ubiquitin substrate profiling

ESC lysate (40 mg) was prepared under denaturing conditions, reduced and alkylated, diluted 4-fold, and then subjected to overnight trypsin digestion. Peptides were acidified with trifluoroacetic acid (TFA), desalted by solid-phase extraction, and lyophilized for 48 h. Dry peptides were resuspended in 1x IAP buffer (Cell Signaling Technologies), clarified by centrifugation, then incubated with anti-KGG coupled resin (Cell Signaling Technologies) for 2 h at 4°C. Beads were washed twice with IAP buffer, four times with water, and eluted twice using 0.15% TFA. Peptide eluates were desalted using STAGE-Tips and analyzed using an Orbitrap-Elite mass spectrometer as described previously (27). Peptide spectral matching was performed using semi-tryptic specificity and ±25 ppm mass precursor tolerance using Mascot (Matrix Science). A target-decoy database comprised mouse proteins (UniProt ver. 2011_12) and common contaminants. Oxidized methionine (+15.9949) and K-GG (+114.0429) were considered as variable modifications, carbamidomethyl cysteine (+57.0214) as a fixed modification, and two missed cleavage events were permitted. Peptide spectral matches were filtered serially using linear discriminant analysis to 5% and 2% FDR at the peptide and protein levels, respectively. Label–free peak areas were determined for confidently identified peptides using the XQuant algorithm and consolidated at the protein level using linear mixed effects modeling as described previously (28).

### Global proteome and phosphoproteome analysis

Cells were lysed in 20 mM HEPES pH 8.0, 9 M urea, and phosphatase inhibitor cocktail (Roche). Lysates were sonicated on a microtip and insoluble material was removed by centrifugation. Approximately 30 mg of total protein per condition was reduced and alkylated in 5 mM DTT and 15 mM iodoacetamide, respectively. Protein mixtures were diluted 4-fold in 20 mM HEPES, pH 8.0 and sequentially digested with LysC (1:50, enzyme:protein, Wako) and trypsin (1:100, enzyme:protein, Thermo Fisher Scientific). Peptide mixtures were purified on SepPak columns (Waters) and lyophilized to dryness. An aliquot of the digested peptides from each condition was processed for global proteome profiling. Briefly, 200 µg of total peptides per condition were labeled with tandem mass tag (TMT10plex and TMT11, Thermo Fisher Scientific). After >98% TMT incorporation rate was confirmed, samples were combined and fractionated by high pH Reverse Phase LC (Agilent Technologies) into 24 fractions for analysis as described previously (29).

From the majority of the sample, phosphopeptides were enriched using TiO_2_ titansphere resin (GL Sciences). Peptides were reconstituted in 50% ACN and 2 M lactic acid buffer, and incubated at 1:5 peptide:titansphere ratio for 2 h at room temperature. The titansphere particles were washed sequentially in 50% ACN/2M lactic acid, 50% ACN/0.1% TFA, and 25% ACN/0.1% TFA. Phosphopeptides were eluted with 50 mM K_2_HPO_4_ pH 10 buffer and lyophilized. Enriched peptides were labeled with TMT10plex and TMT11 reagents (Thermo Fisher Scientific). Samples were combined and processed for phosphotyrosine (pY) enrichment using p100 PTMScan reagent (Cell Signaling Technologies). The flow-through fraction (pST) was subject to high pH reverse phase fractionation into 24 fractions for analysis as described previously (29).

### Liquid chromatography and tandem mass spectrometry

TMT-labeled peptides for global proteome, KGG, pST and pY profiling were desalted on C18 stage tip, dried by vacuum centrifugation, and reconstituted in 2% ACN/0.1% FA for analysis. All samples were analyzed by liquid chromatography coupled to tandem mass spectrometry (LC-MS) on Dionex Ultimate 3000 RSLCnano system (Thermo Fisher Scientific) and Orbitrap Fusion Lumos Tribrid MS (Thermo Fisher Scientific). Peptide samples were resolved by 158 min linear gradient of 2% to 30% buffer B (98% ACN/0.1% FA) in buffer A (2% ACN/0.1% FA) on 100 µm ID PicoFrit column packed with 1.7 µm Acquity BEH (New Objective) at a flow rate of 450 nL/min. Total run length including injection, gradient, column washing and re-equilibraton was 180 min. Multi-Notch MS3-based TMT method was used for sample analysis where the duty cycle involves collecting: 1) an MS1 scan in the Orbitrap at 120,000 resolution across 350-1350 m/z range with automatic gain control (AGC) target of 1.0e6, 50 ms maximum injection time; 2) data dependent ion trap MS2 scans on the top 10 peptides with CID activation, 0.5 m/z isolation window in quadrupole, turbo scan rate, 2.0e4 AGC target, 100 ms maximum injection time; and 3) Orbitrap SPS-MS3 scans of 8 MS2 fragment ions, with isolation widths of 2 m/z using isolation waveforms with multiple frequency notches as described previously (30). MS3 precursor ions were fragmented by high energy collision-induced dissociation and analyzed by Orbitrap at 50,000 resolution, AGC target 2.5e5, 150 ms maximum injection time.

### Data analysis for global and phospho-proteome multiplexed proteomics

MS/MS spectra were searched using the Mascot algorithm (Matrix Sciences) against a concatenated target-decoy database comprised of the UniProt Mus musculus protein sequences (downloaded June 2016), known contaminants, and the reversed versions of each sequence. A 50 ppm precursor ion mass tolerance and 0.8 Da fragment ion tolerant were used with tryptic specificity, and up to 2 missed cleavages permitted. Fixed modifications were considered for carbamidomethylated cysteine residues (+57.0215 Da) and the TMT modification of the N-terminus and K residues (+229.1629 Da). Variable modifications were permitted for methionine oxidation (+15.9949 Da), phosphorylation on S/T/Y (+79.9663 Da) and TMT modification of Y (+229.1629 Da). Search results were filtered to 1% FDR at the peptide level and 2% FDR at the protein level using in house tools as described previously (28). Phospho-sites on peptides were localized with Ascore (31) and all peptides spanning phospho-sites were grouped using their residue and position nomenclature prior to modeling. MS3 based TMT quantification was performed using our in-house Mojave module (32), filtering out TMT peaks in MS3 scans whose reporter ion intensity sum < 30,000, across all 11 channels. For each peptide, the respective reporter ion intensities were summed across PSMs. Sequences <7 residues were further removed due to ambiguity in peptide to protein mapping. Then, for each protein or phospho-site, a model was fitted in MSstats v3.7.1 (33) using Tukey Median Polish summary on all quantified peptides across replicates with imputation of missing values below a censoring threshold of 2^8^. Within MSstats, the model estimated fold change and statistical significance were computed for all compared treatment groups. Significantly altered proteins or phospho-sites were determined by setting a threshold of |Log2(fold-change)| > 1 and p-value < 0.05. The subset of significantly altered phospho-sites, unaltered at the protein level, was then annotated and tested for over-represented biological annotations using the MsigDB collection and GSEA (34). Significant annotations were defined by an enrichment q-value < 0.05.

### Lentivirus and PEG10 VLP generation

HEK293T cells (10^7^) plated on gelatin-coated 10 cm dishes were transfected with 5 µg pGCMV-MCS-IRES-eGFP (Genentech), Delta 8.9 or Delta 8.9-PEG10 (PEG10 residues 1-377 were cloned into the p24 region of the Delta 8.9 vector), and VSV-G at a molar ratio of 1:2.3:0.2 using Lipofectamine 2000 (Thermo Fisher). Media was replenished 6 h post-transfection and the supernatant collected after a further 40 h. Lentivirus particles or VLPs were isolated using a Lenti-X Concentrator (321231, Clontech).

### Protease protection assay

Lentiviruses and PEG10 VLPs were resuspended in PBS. Half of the sample was incubated with 1% Triton X-100, 0.5% TriButyl, and 0.2% SDS, and sonicated briefly to permeabilize the lipid bilayer. 1 µg trypsin (V511A, Promega) was added to both samples (−/+ Triton) and incubated for 0, 30, 60, and 120 min on ice. Trypsinization was stopped by adding 5 mM PMSF. Samples were boiled for 10 min in NuPAGE LDS Sample Buffer (NP0008, Thermo Fisher) and NuPAGE Sample Reducing Agent (NP0004, Thermo Fisher).

### Vesicle isolation

Culture supernatants were centrifuged at 1500 x g, then 10,000 x g, and then filtered through a 0.22 µm vacuum filter unit. Further centrifugation at 100,000 x g for 2 h yielded a vesicle-containing pellet. For sucrose gradient purification, the pellet was resuspended in 60% sucrose in 20 mM Tris-HCl pH 7.4 and 0.85% NaCl. This suspension was layered with 40% and 20% sucrose in 20 mM Tris-HCl pH 7.4 and 0.85% NaCl, and centrifuged at 150,000 x g for 14 h.

### Microscopy

For immuno-electron microscopy, vesicles were adsorbed onto formvar/carbon-coated grids for 20 min, and then fixed and permeabilized for 30 min with 4% paraformaldehyde (PFA) in PBS containing 5% Triton X-100. Samples were quenched in PBS/glycine, rinsed in PBS, and labeled for 1 h with 20 µg/mL anti-PEG10 antibody diluted in EM blocking solution (EMS, Aurion). Samples were washed in PBS for 15 min, and then incubated for 1 h with goat anti-rat 6 nm gold conjugate (Jackson ImmunoResearch) diluted 1/10 in EM blocking solution. Samples were washed in PBS for 10 min, rinsed in water, and stained with 2% ammonium molybdate for 30 sec. Samples were imaged with JEOL JEM-1400 TEM, Ultrascan 1000 CCD camera. For negative stain microscopy, samples were adsorbed to grids for 20 min, rinsed with water, and stained with 1% uranyl acetate for 60 sec.

### Immunohistochemistry

Sections of formalin-fixed, paraffin-embedded C57BL/6J mouse tissues (5 µm) were baked for 20 min at 70°C. Deparaffinization and hydration was performed in a Leica Autostainer XL (Leica Biosystems). Sections were quenched in 3% H_2_O_2_ for 4 min, blocked using a ScyTek Blocking Kit (BBK120, ScyTek Laboratories), and then incubated in 10% donkey serum in 3% BSA/PBS for 30 min. Labeling was for 1 h with 5 µg/ml anti-PEG10 antibody or IgG2b isotype control (553986, BD Pharmingen) in PBS containing 10% donkey serum and 3% BSA, followed by 30 min with 5 µg/ml biotinylated donkey anti-rat antibody (712-065-153, Jackson Laboratories). Sections were treated with Vectastain ABC Elite Reagent (PK-6100, Vector Labs) and then incubated with Pierce Metal Enhanced DAB (3406, Thermo Fisher) for 5 min. Counterstaining was with Mayers filtered hematoxylin.

### Immunofluorescence labeling

Cells were fixed with 8% PFA, blocked for 1 h with 10% donkey serum, 2% BSA, and 0.2% saponin in PBS, incubated overnight at 4°C in 5 µg/ml anti-PEG10 antibody in blocking buffer, and then for 1 h at room temperature in 2 µg/mL Alexa 488-conjugated donkey anti-rat antibody (A-21208, Thermo Fisher). Slides were mounted with ProLong Gold containing DAPI (P36931, Thermo Fisher). Confocal images were captured with a LEICA SP5 laser-scanning confocal microscope.

### RNA sequencing

Total RNA was extracted using a RNeasy Mini Kit (Qiagen), quantified in a NanoDrop 8000 (ThermoFisher), and assessed for integrity using both 2100 5 Bioanalyzer and 2200 TapeStation (Agilent). Libraries were prepared from 1 µg of total RNA with TruSeq RNA Sample Preparation Kit v2 (Illumina). Library size was confirmed using 2200 TapeStation and High Sensitivity D1K screen tape (Agilent), and concentration was determined by Library quantification kit (KAPA). Libraries were multiplexed five per lane and sequenced in a HiSeq2500 (Illumina) to generate 50 million paired end 75 bp reads. Filtering of fastq sequence files removed poor quality reads (read length < 18 or > 30% of cycles with Phred score < 23). Raw FASTQ reads were aligned to the mouse reference genome (GRCm38-mm10) using GSNAP (with parameters -M 2 -n 10 -B 2 -i 1 -N 1 -w 200000 -E 1 --pairmax-rna=200000 --clip-overlap). Reads were filtered to include only the uniquely mapped reads. Differential expression analysis was performed using the voom/limma R package (35). Genes were considered differentially expressed if the log2 fold change was > 1 or < −1, and adjusted p-value < 0.05. Pathway analysis was performed with the R package EGSEA (36).

### eCLIP

eCLIP was performed by Eclipse BioInnovations Inc (San Diego) essentially as described (37). In brief, 2×10^7^ TSCs were UV (254 nm)-crosslinked at 0.4 J/cm^2^, snap frozen, and then lysed prior to treatment with RNase I. PEG10 was immunoprecipitated with anti-PEG10 antibody pre-coupled to sheep anti-mouse Dynabeads (Thermo Fisher) overnight at 4°C. 20% of the sample was used for western blotting, 2% was used as a paired input, and the remainder was processed for eCLIP. Raw FASTQ files were trimmed of 3’ barcodes and adapter sequences. Reads were aligned to the mouse reference genome (GRCm38-mm10) using STAR {Dobin, 2013 #1496} followed by removal of PCR duplicates and reads mapping to repetitive elements in the genome. MACS2 {Zhang, 2008 #1426} was used to call peaks in wild-type samples using PEG10-deficient samples as controls. In Fig 7B, each PEG10 bound gene was divided into 100 bins between the Transcription Start Site (TSS) and the Transcription End Site (TES), and into 20 bins upstream and downstream of TSS and TES, respectively. For each bin, count of 5’-end of each eCLIP read was calculated and summed across all genes to create the average profile per sample. All samples were normalized to sequencing depth.

### Raw data deposition

All mass spectrometry raw files were uploaded on the Massive data repository and can be downloaded from ftp://MSV000083229@massive.ucsd.edu (login: MSV000083229_reviewer, password: a). eCLIP and RNAseq data have been deposited to the Gene Expression Omnibus (GEO accession number GSE122217). Data can be accessed using the token “kpmneycydrcvdat”.

## RESULTS AND DISCUSSION

### PEG10 is regulated by USP9X

We identified PEG10 while looking for substrates of the deubiquitinating enzyme USP9X in mouse ESCs. USP9X was affinity purified from *Usp9x*^*3xf/y*^ ESCs, which had a 3xFLAG epitope tag inserted in frame with the N-terminus of USP9X. Co-immunoprecipitating proteins captured with anti-FLAG antibody and identified by mass spectrometry included PEG10 (Fig 1A). We confirmed that both PEG10-RF1 and PEG10-RF1/2 also interacted with untagged USP9X using *Peg10*^*+/3xf*^ *Usp9x Rosa26*^*+/CreERT2*^ ESCs, which delete *Usp9x* in response to 4-hydroxytamoxifen and express PEG10 proteins tagged at the N-terminus with 3xFLAG. Importantly, a USP9X antibody co-immunoprecipitated 3xFLAG.PEG10-RF1 and 3xFLAG.PEG10-RF1/2 only when the ESCs expressed USP9X (Fig 1B). Ubiquitin-substrate profiling by K-ε-GG mass spectrometry (38) revealed PEG10 as a putative substrate of USP9X because *Usp9x^C1599A/y^* ESCs expressing catalytically inactive USP9X C1599A (Fig 1C) contained more ubiquitinated PEG10 than wild-type ESCs (Fig 1, D-F). PEG10 deubiquitination by USP9X appeared to stabilize PEG10-RF1 specifically because USP9X-deficient ESCs contained less PEG10-RF1 than control ESCs, whereas the amount of PEG10-RF1/2 was largely unaltered (Fig 1, B and G). Consistent with ubiquitinated PEG10-RF1 being targeted for proteasomal degradation, the proteasome inhibitor MG-132 increased the amount of PEG10-RF1 in USP9X-deficient ESCs, but had little impact on PEG10-RF1/2 (Fig 1G). PEG10-RF1 has six unique residues that are located in the frameshift region and are not found in PEG10-RF1/2 (12), so these may contribute to a degron motif that specifies its proteasomal degradation. We did not detect any ubiquitination sites within the frameshift region, but USP9X suppressed ubiquitination of two adjacent lysines (K365, K362). Whether mutation of these lysines impacts PEG10-RF1 stability is an area of future investigation.

**Fig 1.**
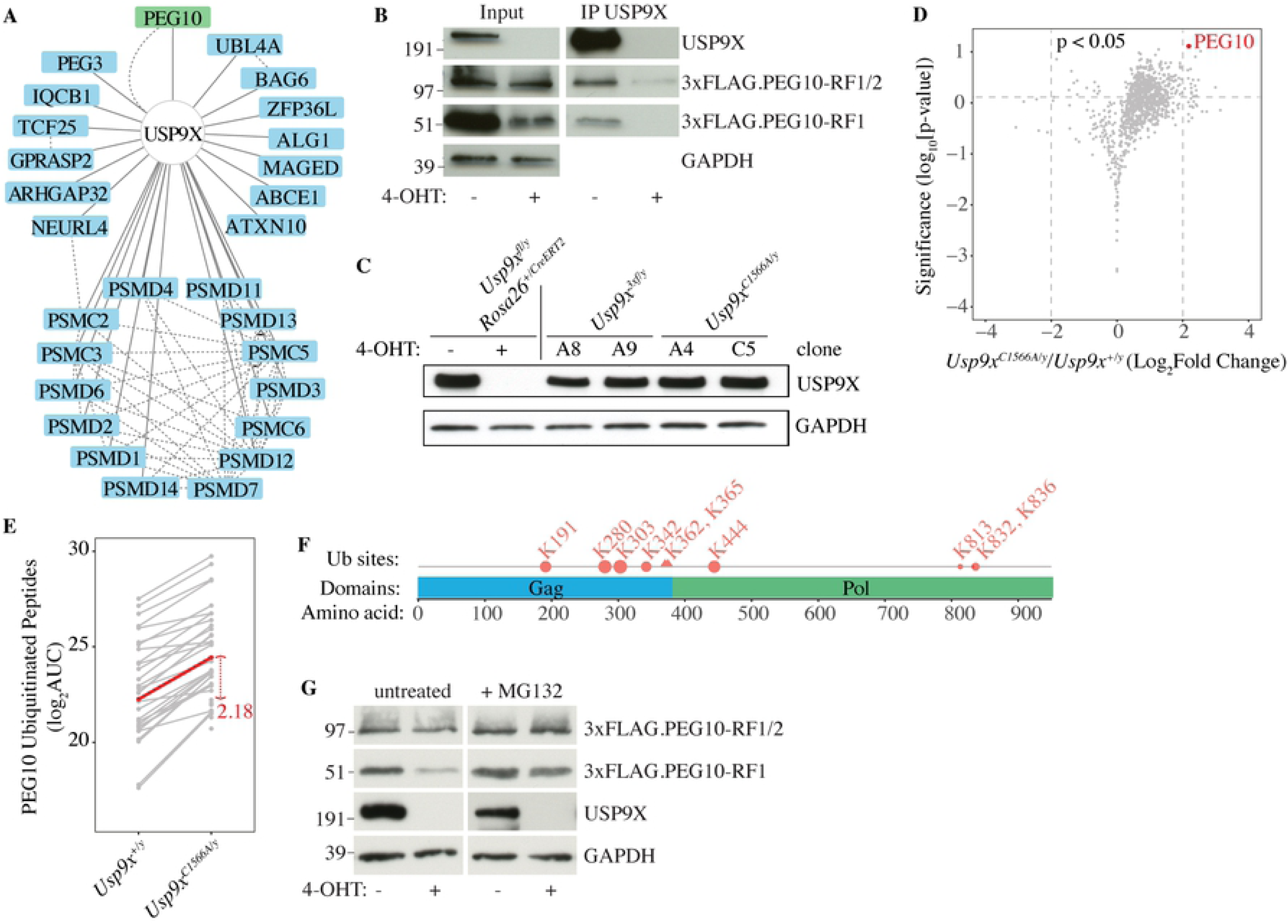
PEG10 is a substrate of USP9X. (A) USP9X interaction network showing all high confidence binding partners connected by solid lines (Saint > 0.9, FDR < 0.05, Avg. Psms > 10). Dotted lines indicate interactions reported in the BioPlex network (50, 51). Results are representative of 3 independent experiments. (B) Western blots of *Peg10*^*3xf/+*^ *Usp9X*^*fl/y*^ *Rosa26*.CreER^T2^ ESCs after immunoprecipitation (IP) of USP9X. Epitope-tagged PEG10 was detected with anti-FLAG antibody. Where indicated, ESCs were treated with 4-hydroxytamoxifen (4-OHT) to delete *Usp9x*. Results are representative of 2 independent experiments. (C) Western blots of ESCs. (D) Geyser plot showing proteins differentially ubiquitinated in *Usp9x*^*C1566A/y*^ ESCs versus control *Usp9x*^*+/y*^ ESCs (p < 0.05, log_2_ fold change > 2). Results are representative of 2 independent experiments. (E) Line plot showing the fold-change in abundance of all ubiquitinated PEG10 peptides between *Usp9x*^*+/y*^ and *Usp9x*^*C1566A/y*^ ESCs. Each grey line corresponds to a unique KGG peptide spectral match peptide. The red line indicates a model-based protein abundance estimate. AUC, area under the curve. (F) Diagram indicating the position of the USP9X-regulated ubiquitination (Ub) sites in PEG10. Triangles depict ubiquitination sites identified only in *Usp9x*^*C1566A/y*^ ESCs. Circles depict sites found in *Usp9x*^*+/y*^ and *Usp9x*^*C1566A/y*^ ESCs. Circle size indicates the extent to which ubiquitination was increased by USP9X inactivation. (G) Western blots of *Peg10* ^*3xf/+*^ *Usp9x*^*fl/y*^ *Rosa26*.CreER^T2^ ESCs. Where indicated, cells were treated with 10 μM MG-132 for 4 h. Results are representative of 2 independent experiments.

### The PEG10 Gag domain forms virus-like particles

The biochemical roles of PEG10 are unknown. The Gag polyproteins of Ty3 retrotransposons have been shown to assemble virus-like particles (VLPs) (39), so we explored whether the conserved PEG10 Gag domain could substitute for the p24 HIV-1 Gag protein in a lentiviral packaging system. HEK293T cells were co-transfected with VSV-G envelope protein and either HIV-1 Gag-Pol or a hybrid construct encoding PEG10-RF1 and HIV-1 Pol. Both p24 and PEG10-RF1 were detected in the culture supernatant by western blotting (Fig 2A), suggesting that both Gags assembled VLPs. Sucrose gradient centrifugation suggested the putative PEG10-RF1 VLPs were denser than the HIV-1 p24 VLPs (Fig 2B). VLPs that bud from the cell are encapsulated in a lipid bilayer that protects Gag proteins from digestion with trypsin. Accordingly, PEG10-RF1 in culture supernatants was only cleaved by trypsin in the presence of the bilayer-disrupting detergent Triton X-100 (Fig 2C). Indeed, lipid bilayer-encapsulated PEG10-RF1 VLPs were detected by negative stain electron microscopy (Fig. 2D).

**Fig 2.**
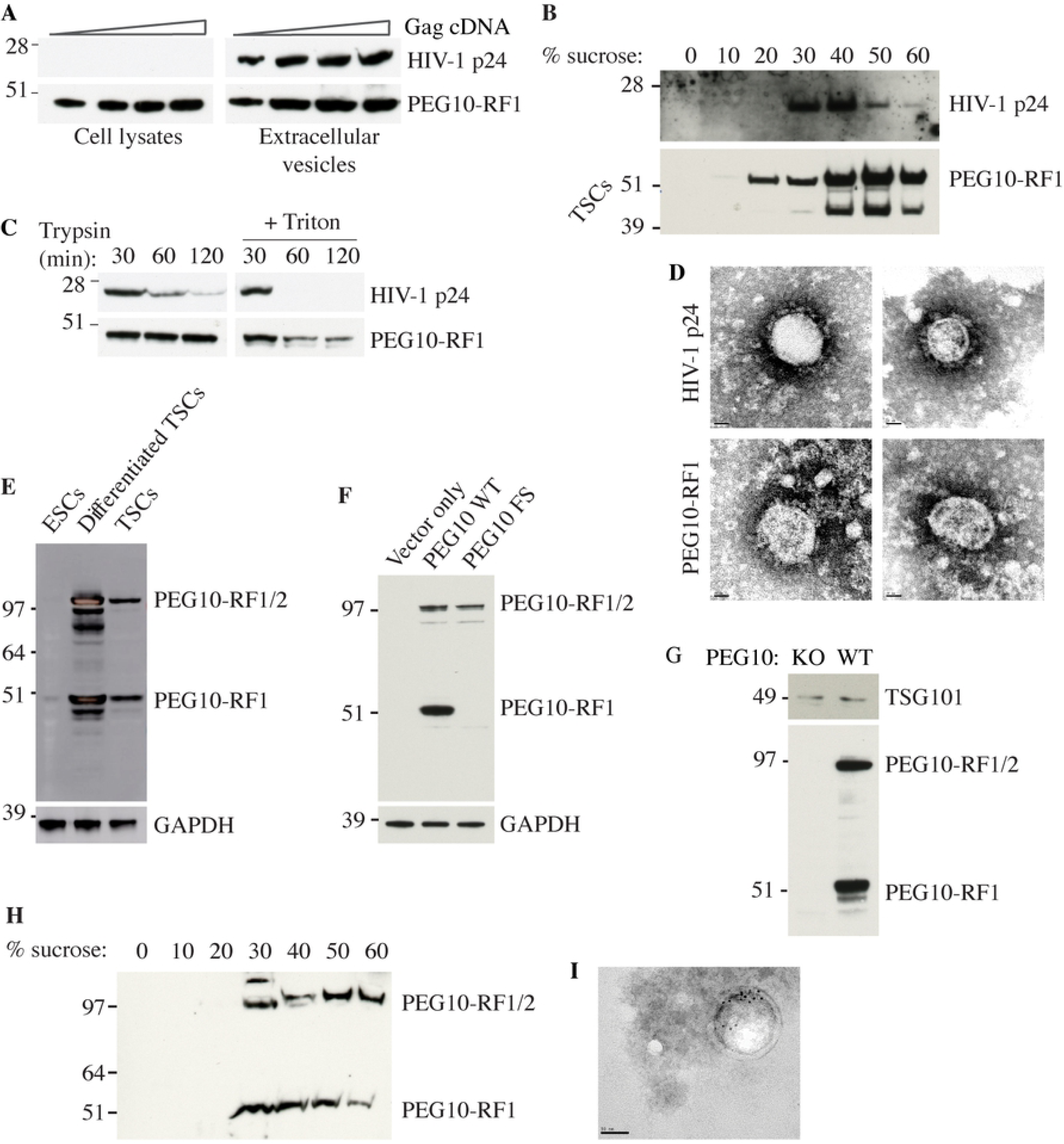
The PEG10 Gag domain generates virus-like particles. (A) Western blots of HEK293T cells or extracellular vesicles recovered from the culture medium after transfection with VSV-G and increasing amounts of Gag-Pol cDNA. Results are representative of 2 independent experiments. (B) Western blots of extracellular vesicles in panel A after sucrose gradient density centrifugation. The upper blot shows cells transfected with HIV-1 Gag, whereas the lower blot shows cells transfected with PEG10-RF1. We speculate that the faster migrating PEG10-RF1 species is a processed form of the protein. Results are representative of 2 independent experiments. (C) Western blots of VLPs. Results are representative of 3 independent experiments. (D) Electron micrographs of negatively stained VLPs. Scale bar, 20 nm. Results are representative of 2 independent experiments. (E) Western blots of wild-type ESCs and TSCs. Results are representative of 2 independent experiments. (F) Western blots of HEK293T cells ectopically expressing wild-type (WT) mouse PEG10 or a PEG10 frameshift (FS) mutant that can only make PEG10-RF1/2. Results are representative of 2 independent experiments. (G) Western blots of extracellular vesicles shed from WT or PEG10-deficient (KO) TSCs and enriched by differential centrifugation. Results are representative of 5 independent experiments. (H) Western blots of extracellular vesicles shed from WT TSCs and analyzed by sucrose gradient density centrifugation. Results are representative of 2 independent experiments. (I) Electron micrographs of TSC extracellular vesicles after immuno-gold labeling for PEG10. Scale bar, 50 nm.

Next, we determined if endogenous PEG10 was in vesicles shed from trophoblast stem cells (TSCs), which express even more PEG10 than ESCs (Fig 2E). PEG10 was detected with a monoclonal antibody that was raised against PEG10-RF1, and therefore detects both PEG10-RF1 and PEG10-RF1/2 (Fig 2, E and F). We found that both PEG10-RF1 and PEG10-RF1/2 were present in supernatant fractions enriched for TSC extracellular vesicles (Fig 2, G and H). The exosome marker TSG101 (40) served as a positive control. Electron microscopy and immuno-gold labeling for PEG10 confirmed the presence of PEG10-bearing vesicles (Fig 2I). Collectively, our data suggest that the PEG10 Gag domain supports constitutive VLP assembly.

To investigate the mechanism of PEG10 budding, we affinity purified PEG10 from ESCs and TSCs and then identified co-immunoprecipitating proteins by mass spectrometry (Fig 3A). Consistent with our previous experiments, PEG10 interacted with USP9X and USP9X-associated proteins such as ATXN10, IQCB1, BAG6 and UBL4. Intriguingly, two proteins involved in vesicle fusion, VTI1B and STX8 (41, 42), also coimmunoprecipitated with PEG10 (Fig 3, A and B). It is tempting to speculate that VTI1B and STX8 might regulate the budding of PEG10-containing vesicles.

**Fig 3.**
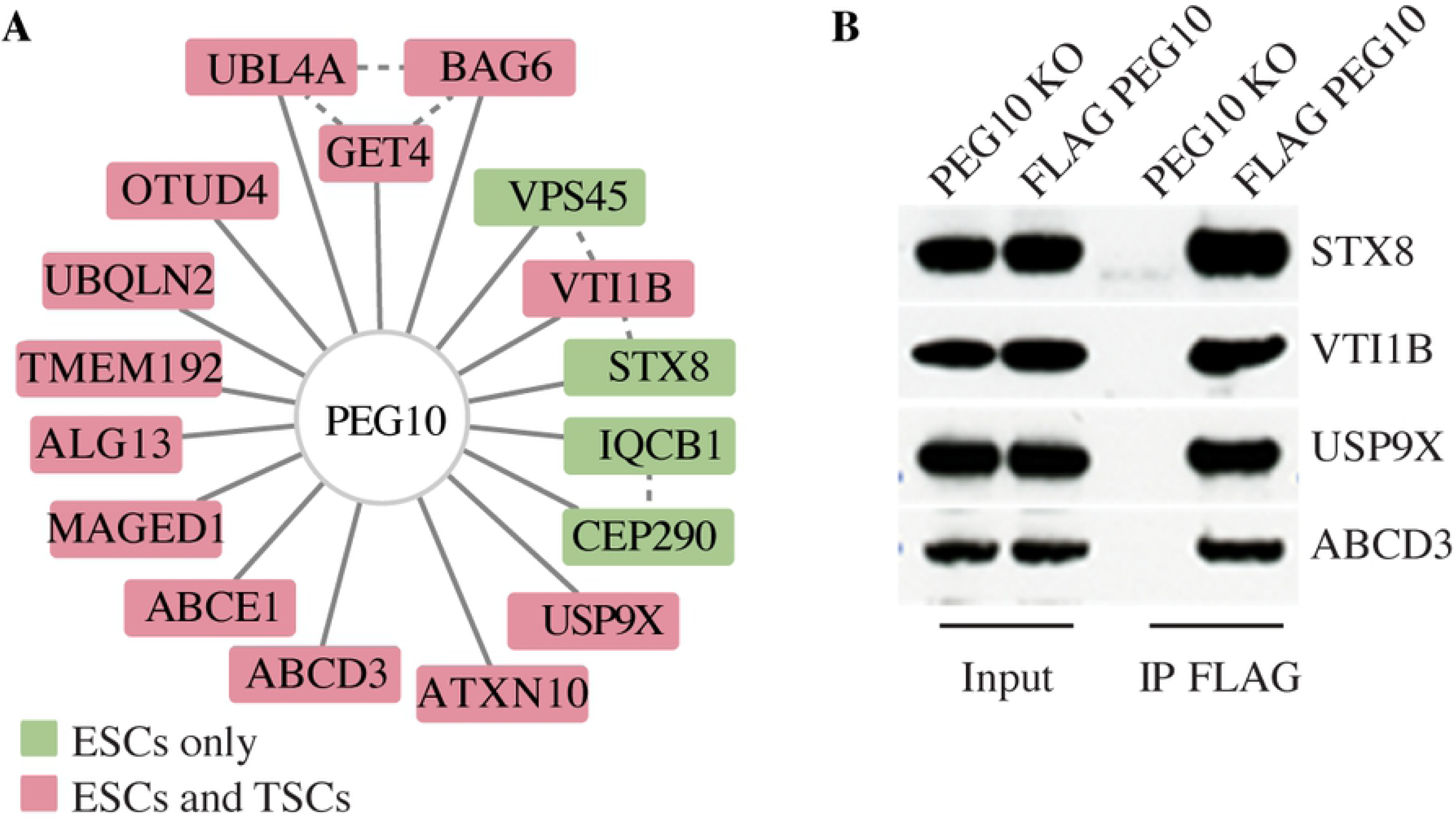
Proteins interacting with PEG10 in ESCs and TSCs. (A) PEG10 interaction network showing all high confidence binding partners connected by solid lines (Saint > 0.9, FDR < 0.05, Avg. Psms > 10). Dotted lines indicate interactions reported in the Bioplex interaction network. Proteins co-immunoprecipitated from both TSCs and ESCs are colored red, and those unique to ESCs are green. Results are representative of 3 independent experiments. (B) Western blots before and after immunoprecipitation (IP) of 3xFLAG.PEG10 from knock-in ESCs, or as a control, PEG10-deficient (KO) ESCs. Results are representative of 2 independent experiments.

### PEG10 is expressed in a subset of adult mouse tissues

PEG10 has been studied largely in developmental settings and in cancer cell lines, so we determined its expression in a broad panel of mouse tissues. PEG10-RF1 and PEG10-RF1/2 were both abundant in adrenal gland, testis and placenta, whereas pituitary, ovary, uterus, white adipose, brain and lung expressed them to a lesser extent (Fig 4A). Additional faster migrating bands detected by the PEG10 antibody may reflect proteolytic processing of PEG10. Immunolabeling of tissue sections revealed that PEG10 was expressed highly in the cortex of the adrenal gland, the pars distalis of the pituitary gland, the sertoli cells of the testis, the hypothalamus, and in the labyrinth and trophoblast layers of the placenta (Fig 4B). Expression of PEG10 in the pituitary gland is interesting given that PEG10-deficient mice generated by tetraploid complementation exhibit severe growth retardation (13). PEG10 appeared largely cytoplasmic in the different mouse tissues. Immunofluorescence staining (Fig 4C) and biochemical fractionation studies (Fig. 4D) indicated that PEG10 in ESCs and TSCs was also mostly cytoplasmic. Both PEG10-RF1 and PEG10-RF1/2 were in the cytoplasm of ESCs, but some PEG10-RF1/2 was nuclear in TSCs (Fig. 4D). Therefore, the cellular localization of PEG10 appears context-dependent.

**Figure 4.**
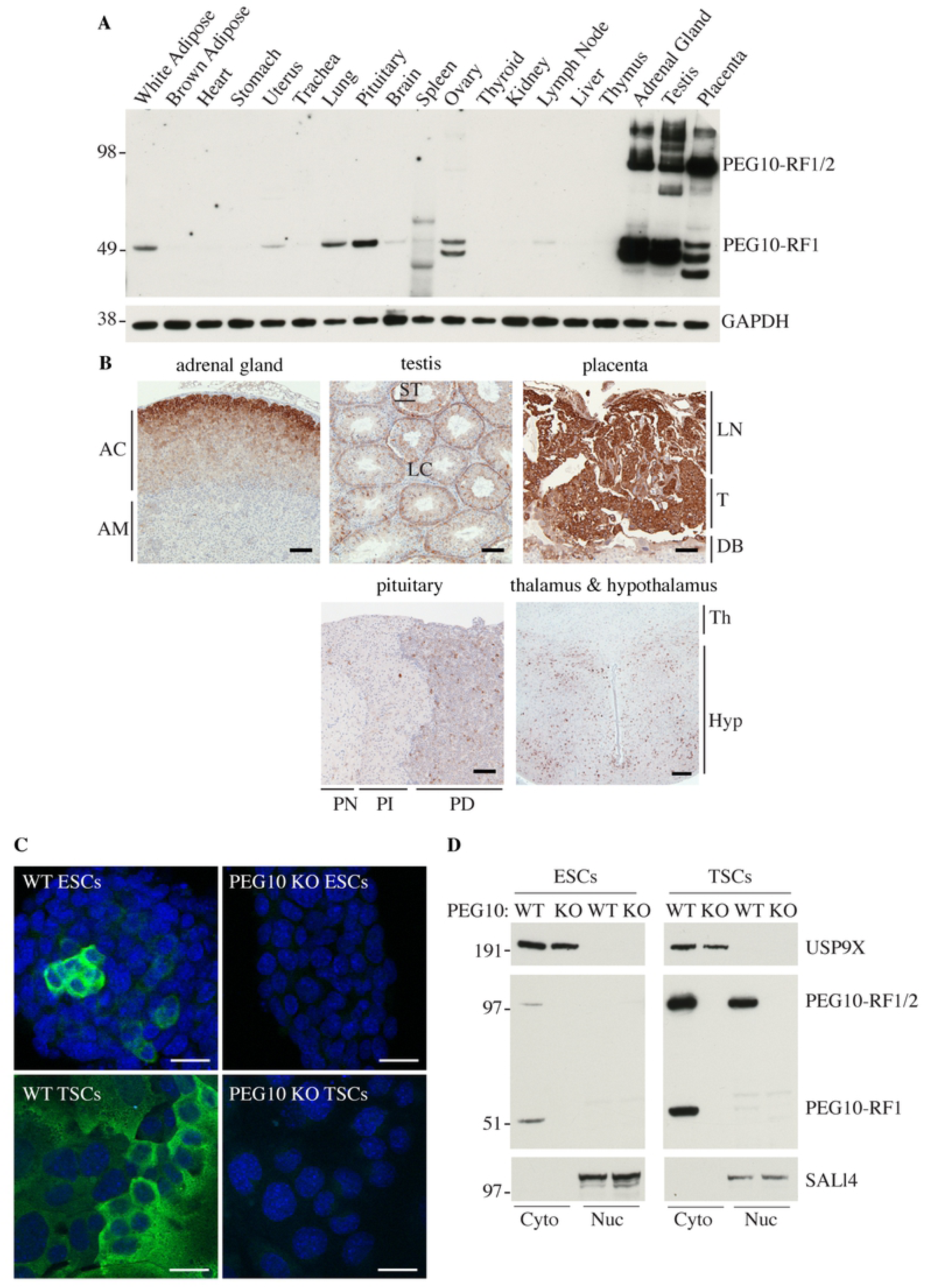
Expression of PEG10 in mouse tissues. (A) Western blots of C57BL/6 mouse tissues. Results are representative of 3 independent experiments. (B) Mouse tissue sections immunolabeled for PEG10 (brown). AC, adrenal cortex. AM, adrenal medulla. LN, Labyrinth. T, Trophoblast. DB, decidua basalis. ST, Sertoli cells. LC, Leydig cells. PN, pars nervosa. PI, pars intermedia. PD, pars distalis. Th, thalamus. Hyp, Hypothalamus. Scale bar, 100 µm. Results are representative of 3 independent experiments. (C) Immunofluorescence staining of PEG10 in ESCs or TSCs. Scale bar, 25 µm. Results are representative of 2 independent experiments. (D) Western blots of ESCs and TSCs that were fractionated into cytoplasmic (cyto) and nuclear (nuc) compartments. Results are representative of 3 independent experiments.

### PEG10 is essential for TSC differentiation

PEG10 is essential for formation of the labyrinth and spongiotrophoblast layers of the placenta (13). TSCs isolated from the early blastocyst can be differentiated into cells of the labyrinth and spongiotrophoblast layers (25), so we utilized this culture system to further explore the function of PEG10. TSCs were maintained in their pluripotent state on CELLstart extracellular matrix in the presence of fibroblast growth factor 4 (Fgf4) and heparin, or they were differentiated in the absence of CELLstart, Fgf4, and heparin in either 20% oxygen, to form the multinucleated syncytiotrophoblasts (SynTs) of the labyrinth, or 2% oxygen, to form the spongiotrophoblasts and trophoblast giant cells (TGCs) of the junctional zone. We noted that expression of PEG10-RF1/2 and PEG10-RF1 increased as TSCs were differentiated in 20% oxygen (Fig 5A). Upregulation of PEG10 was functionally significant because CRISPR-Cas9-mediated deletion of *Peg10* (Fig. 4D) impaired TSC differentiation under both normoxic and hypoxic conditions (Fig 5, B and C). Morphologically, PEG10-deficient cells retained an undifferentiated appearance (Fig 5B), and by RNA sequencing (RNA-seq), they failed to manifest the transcriptional signature of wild-type, differentiated TSCs (Fig 5C; data available in full through the GEO database, accession GSE122217).

**Figure 5.**
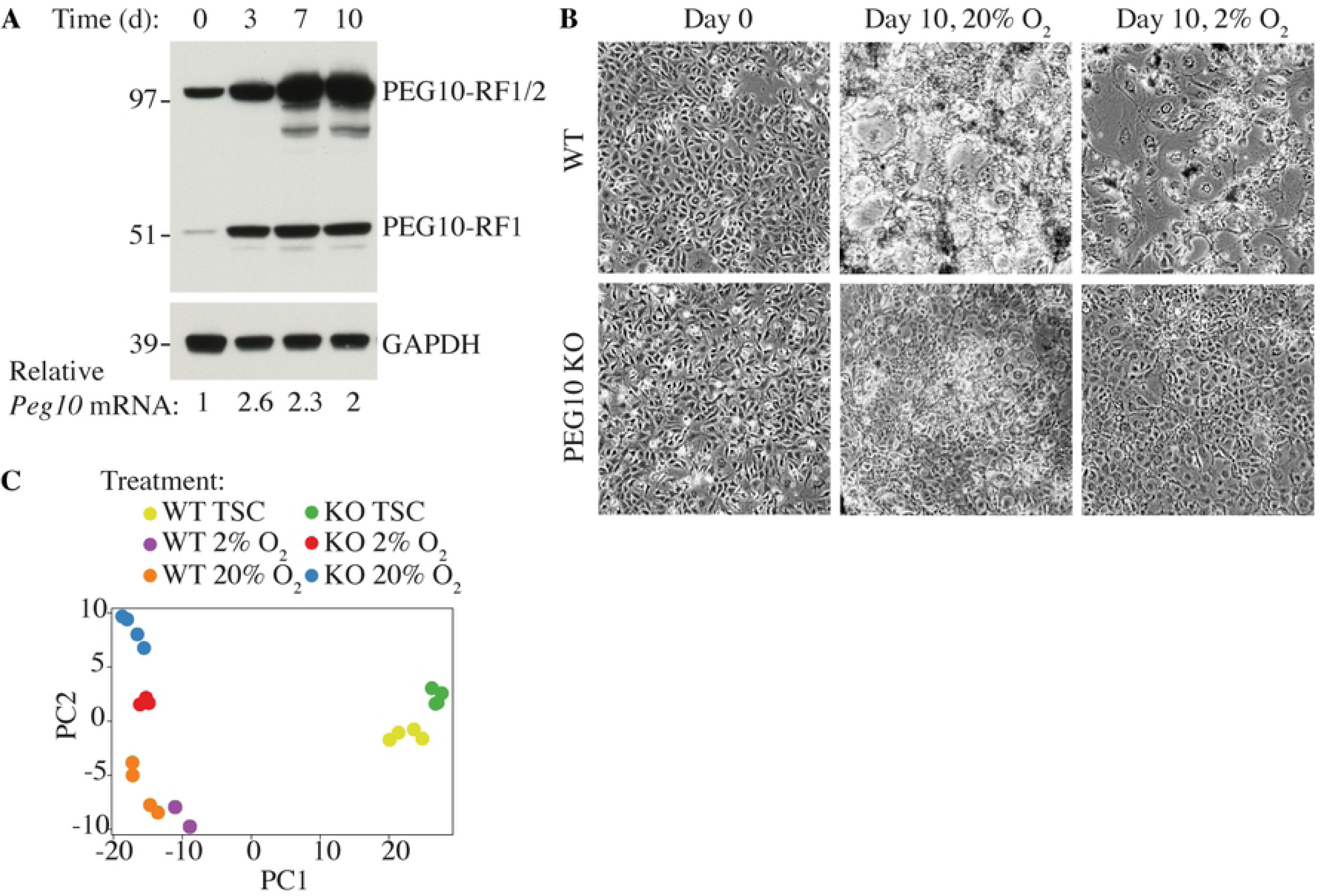
PEG10 regulates the differentiation of TSCs. (A) Western blots of TSCs differentiated in 20% oxygen for the times indicated. *Peg10* mRNA expression was determined by quantitative RT-PCR. (B) Micrographs of wild-type (WT) and PEG10-deficient (KO) TSCs. Results are representative of 5 independent experiments. (C) Principal Component Analysis (PCA) of TSC RNA-seq datasets using log2 RPKM values.

Global proteome and phospho-serine/-threonine/-tyrosine profiling revealed many differences between wild-type and PEG10-deficient TSCs following differentiation (Fig 6, A-D). One particularly interesting difference was the altered phospho-status of key signaling proteins such as MAPK1, MAPK3, MTOR, INSR, and EGFR (Fig 6, B and E). Indeed, enrichment analysis of differentially phosphorylated proteins highlighted aberrant MTOR, Insulin, ErbB, and MAPK signaling (Fig 6F). Phosphoproteome analysis also allowed us to map phospho-sites within PEG10 (Fig 6G).

**Fig 6.**
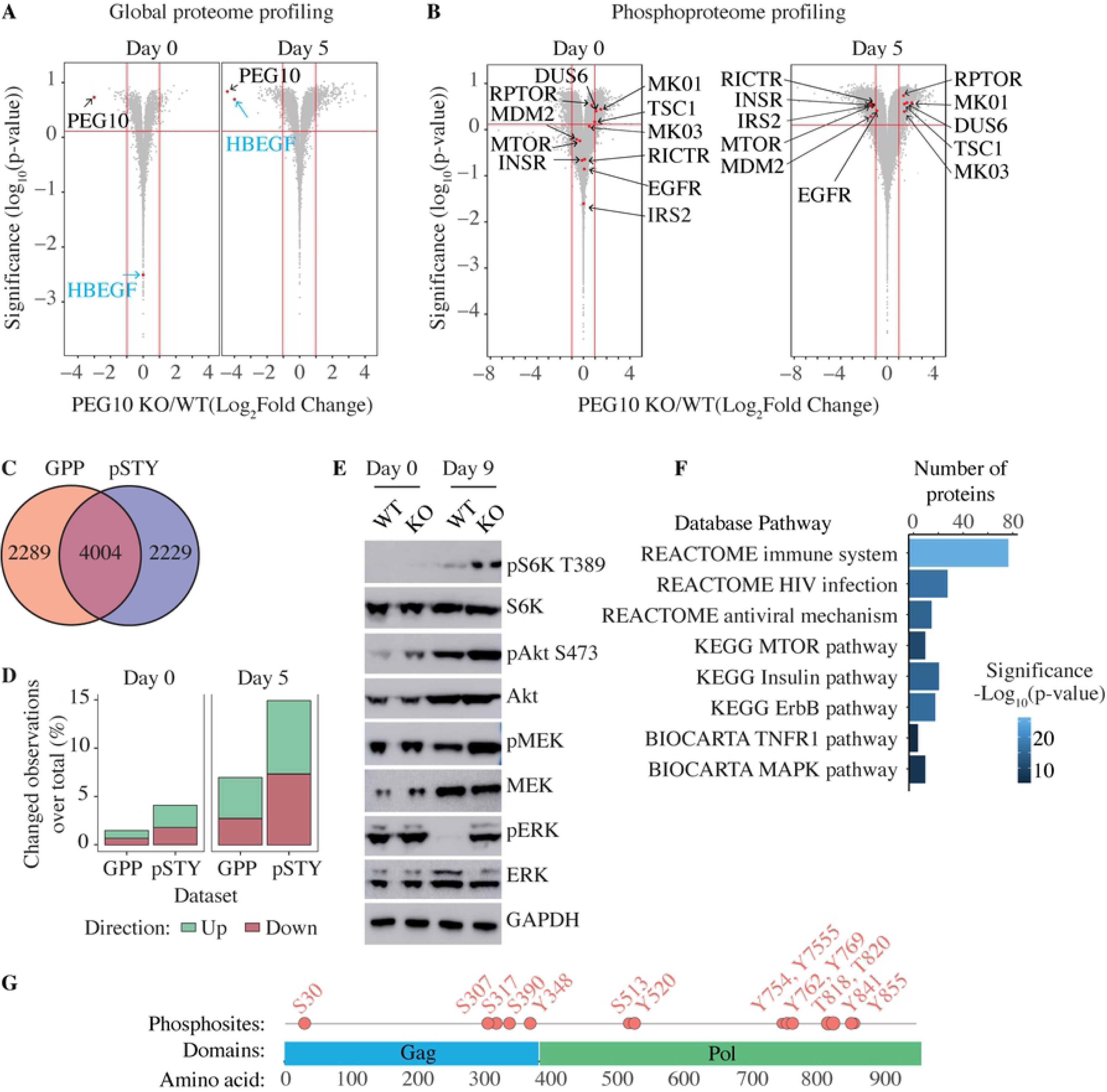
Proteomic analysis of wild-type and PEG10-deficient TSCs. (A and B) Volcano plots of total protein levels (A) or phosphorylation levels at unique phosphosites (B) in wild-type (WT) versus PEG10-deficient (KO) TSCs after 0 and 5 days of differentiation. Results are representative of 3 independent experiments. (C) Venn diagram indicates the total number of proteins identified and quantified by global proteome profiling (GPP) and phosphoproteome profiling (pSTY). (D) Fraction of proteins (GPP) or phosphosites on Ser/Thr/Try (pSTY) with levels changing more than 2-fold (p-value <= 0.05) in PEG10 KO TSCs compared to WT TSCs after 0 and 5 days of differentiation. (E) Western blots of WT and PEG10 KO TSCs after 0 and 9 days of differentiation. (F) Pathways highlighted by the Broad Institute gene set enrichment analysis of proteins with significantly changing phosphorylation levels in PEG10 KO TSCs compared to WT TSCs. (G) Diagram indicates the position of phosphosites in PEG10.

### PEG10 binds to cellular RNAs

The PEG10 Gag domain harbors a conserved zinc finger (ZnF) motif (12) that is reminiscent of the ZnF motifs in orthoretroviral Gags for binding and packaging nucleic acids (43). Therefore, we performed eCLIP-seq (enhanced crosslinking and immunoprecipitation-sequencing) (37) on wild-type and PEG10-deficient TSCs to determine if PEG10 was an RNA binding protein. PEG10 interacted with the 3’ untranslated regions (UTRs) of approximately 840 and 3,680 unique mRNAs after zero and five days of differentiation (normoxic conditions), respectively (Fig 7, A and B; data available in full through the GEO database, accession GSE122217). However, these 3’ UTRs did not have any RNA sequence motifs in common (Fig 7C), which could indicate that RNA secondary structure determines PEG10 binding. PEG10 was bound to its own mRNA (Fig 7D) and to RNAs encoding kinases (MAPK1, MAP2K1, GSK3β, ROCK1, and ROCK2) and regulators of signaling pathways (CMKLR1, TGFa, CBL, GAB1, and HBEGF). Our RNA-seq dataset revealed that a large number of PEG10-bound transcripts, including *Hbegf* (heparin-binding EGF-like growth factor), showed increased expression in wild-type TSCs after differentiation but did not increase in PEG10-deficient TSCs (Fig 7, D and E). HBEGF protein was also less abundant in PEG10-deficient TSCs than in WT TSCs after 5 days of differentiation (Fig 6A). *Hbegf* stood out in this list of genes because of its important role in placentation (44). Therefore, RNA binding by PEG10 appears to promote the expression of genes necessary for differentiation of the placental lineages.

**Fig 7.**
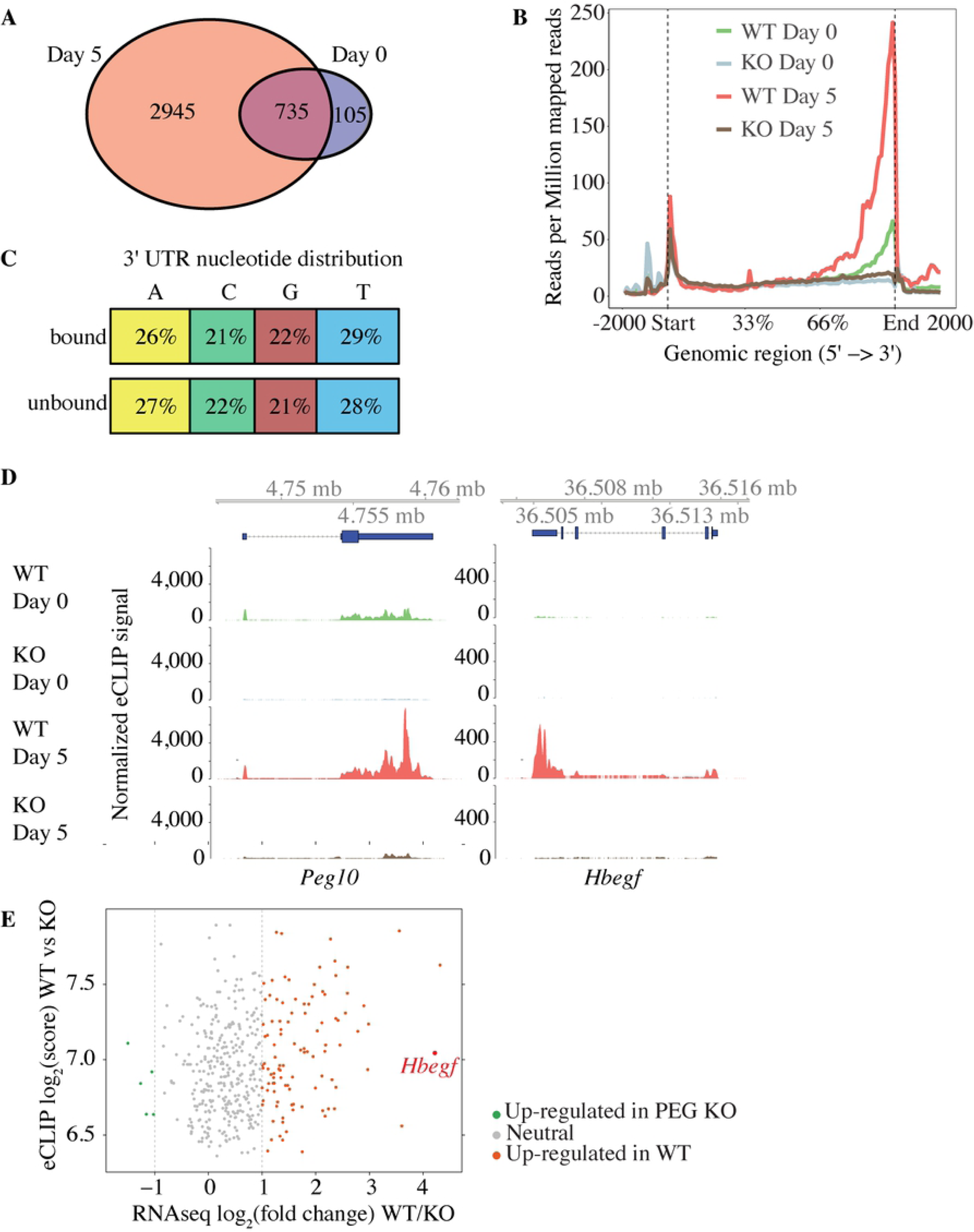
PEG10 binds to RNA. (A) Venn diagram showing RNA transcripts bound to PEG10 after 0 and 5 days of differentiation under normoxic conditions. (B) Graph indicates the distribution of 5’-end eCLIP reads after immunoprecipitating PEG10 from wild-type (WT) and PEG10-deficient (KO) TSCs. Reads are normalized to sequencing depth. (C) Diagram indicates the percentage distribution of nucleotides in PEG10-bound and unbound 3’ UTRs. (D) eCLIP-seq data from WT and PEG10 KO TSCs. Blue boxes at the top indicate the location of the genes. (E) Scatter plot comparing RNA-seq and eCLIP peak scores from WT and PEG10 KO TSCs. Genes are shaded red if they are up-regulated in WT TSCs by RNA-seq (log2 fold change >1 and p-value <0.05), grey if they are unchanged (log2 fold change <1 or >-1), and green if they are down-regulated in WT TSCs (log2 fold change <-1 and p-value <0.05). Results are representative of 2 independent experiments.

In summary, despite its domestication, *Peg10* maintains several hallmarks of retroviral and retrotransposon Gag proteins. The Gag domain of PEG10 supports the budding of virus-like particles, which are released from the cell and can be recovered from exosome preparations. ARC (activity regulated cytoskeletal-associated protein) is another retroviral-like Gag protein that drives the budding of extracellular vesicles (46, 47). ARC, like PEG10, binds to its own mRNA. The release and uptake of *Arc*-containing VLPs by neurons is considered a mechanism of intercellular communication (46, 47). Additional work is needed to determine if PEG10 fulfils a similar function.

## ACKNOWLEDGMENTS

PTMscan® is performed at Genentech under limited license from Cell Signaling Technologies (Danvers, MA). We thank Ben Haley for reagents; Woong Kim and Daisy Bustos for technical assistance.

